## Supplementary material for "Spatial distribution of private gene mutations in clear cell renal cell carcinoma": Additional_File_1.pdf

* Shared first author

### Additional File 1

Supplementary Figures S1-S3

Supplementary Tables S1-S3

**Supplementary Figure S1.** Euclidean distances between samples of each patient. Each sample was sequenced twice producing both single and paired-end reads. Raw count data were adjusted using the variance stabilizing transformation function from DESeq2 (1). Euclidean distances where then calculated. All matrices can be found in Additional File 2.

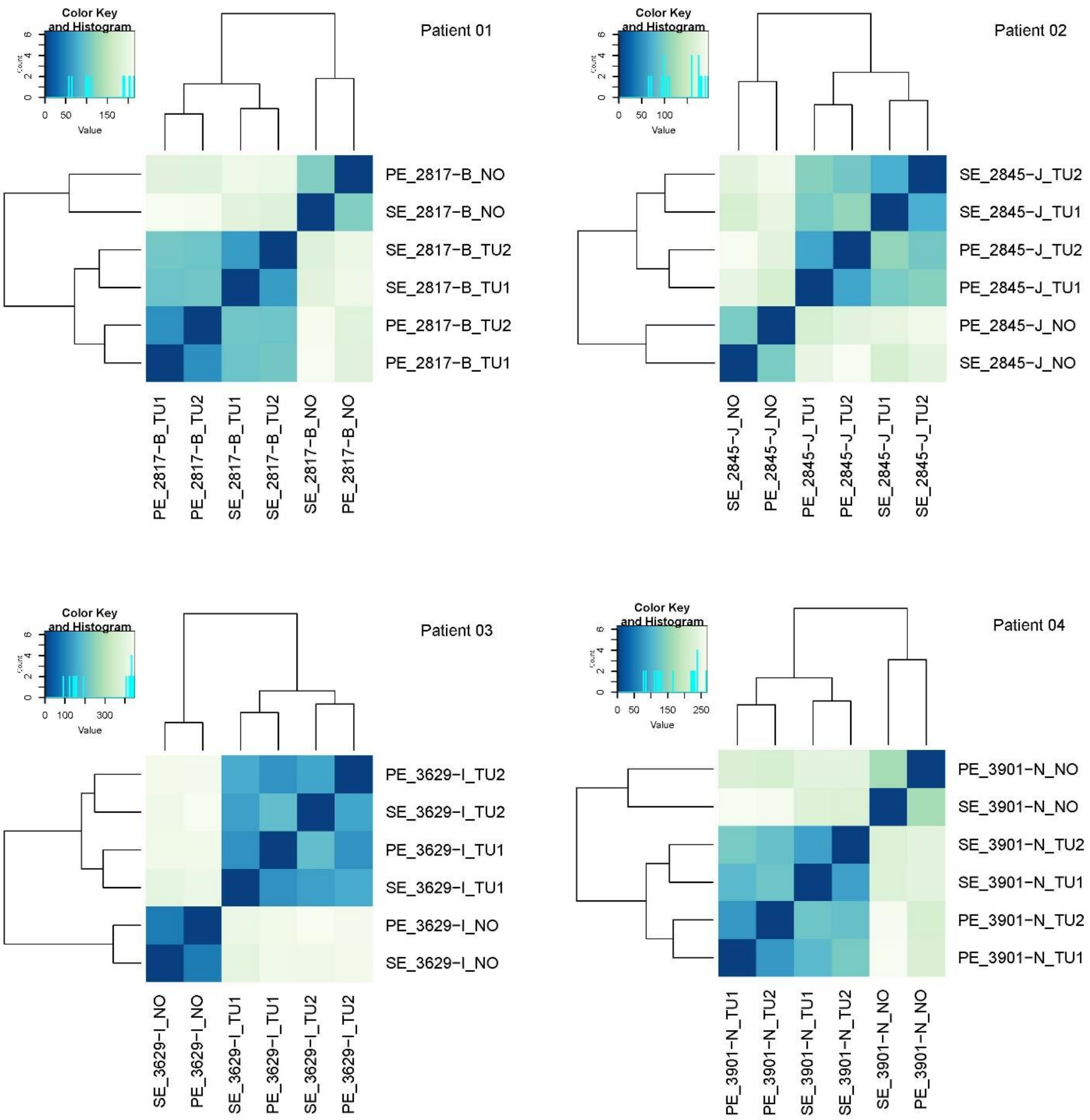

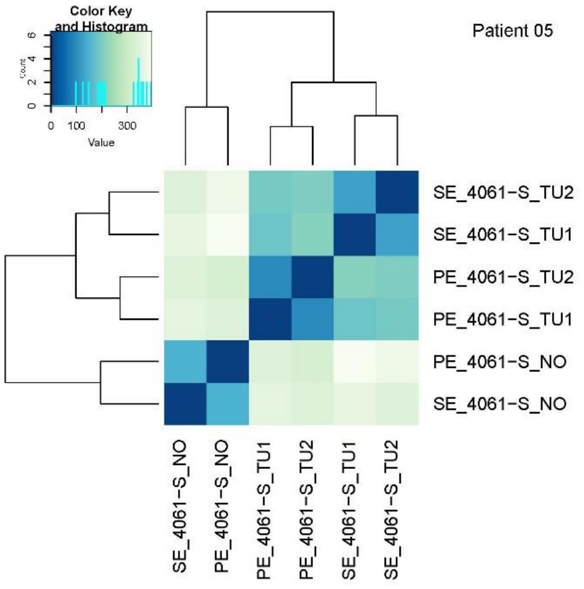
Color Key
and Histogram
Value
Patient 05
SE_4061-S_TU2
SE_4061-S_TU1
PE_4061-S_TU2
PE_4061-S_TU1
PE_4061-S_NO
SE_4061-S_NO
SE_4061-S_NO
PE_4061-S_NO
PE_4061-S_TU1
PE_4061-S_TU2
SE_4061-S_TU1
SE_4061-S_TU2

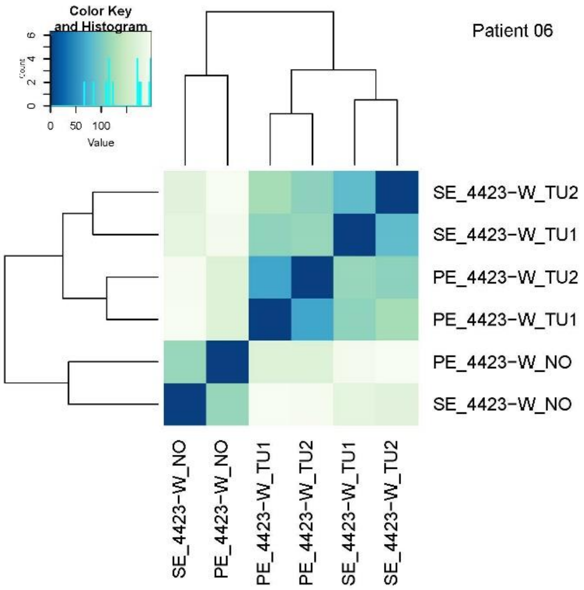
Color Key
and Histogram
Value
Patient 06
SE_4423-W_TU2
SE_4423-W_TU1
PE_4423-W_TU2
PE_4423-W_TU1
PE_4423-W_NO
SE_4423-W_NO
SE_4423-W_NO
PE_4423-W_NO
PE_4423-W_TU1
PE_4423-W_TU2
SE_4423-W_TU1
SE_4423-W_TU2

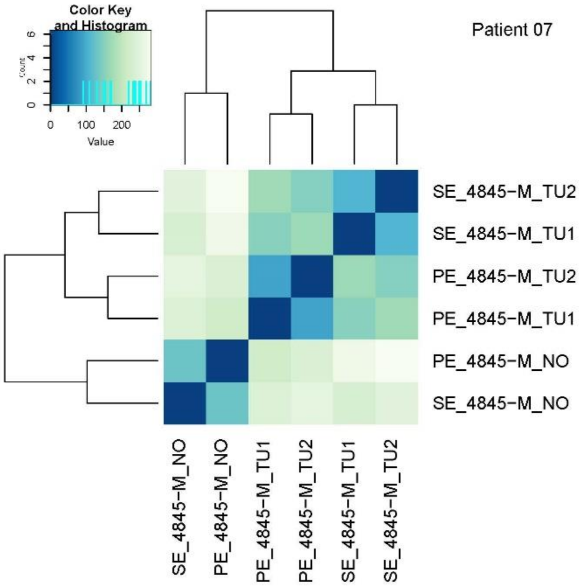
Color Key
and Histogram
Value
Patient 07
SE_4845-M_TU2
SE_4845-M_TU1
PE_4845-M_TU2
PE_4845-M_TU1
PE_4845-M_NO
SE_4845-M_NO
SE_4845-M_NO
PE_4845-M_NO
PE_4845-M_TU1
PE_4845-M_TU2
SE_4845-M_TU1
SE_4845-M_TU2

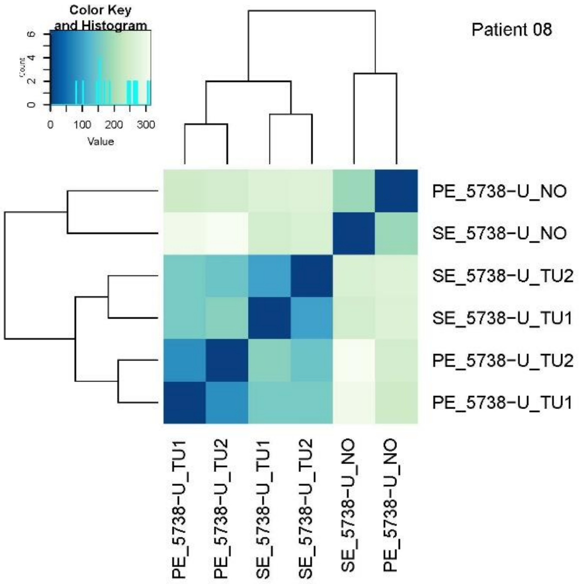
Color Key
and Histogram
Value
Patient 08
PE_5738-U_NO
SE_5738-U_NO
SE_5738-U_TU2
SE_5738-U_TU1
PE_5738-U_TU2
PE_5738-U_TU1
PE_5738-U_TU1
PE_5738-U_TU2
SE_5738-U_TU1
SE_5738-U_TU2
SE_5738-U_NO
PE_5738-U_NO

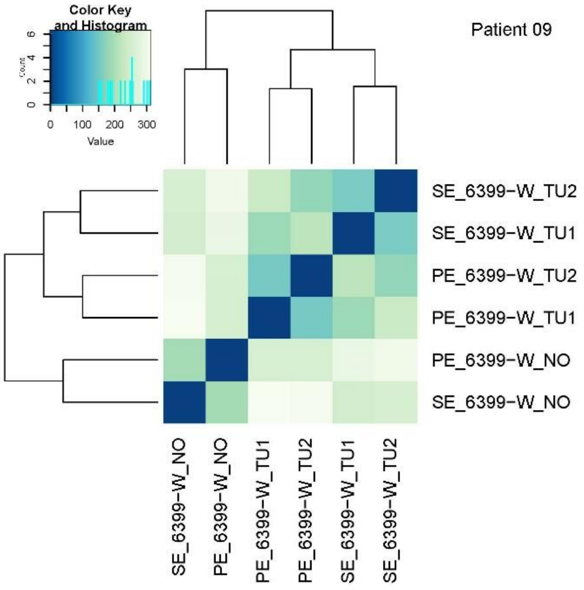
Color Key
and Histogram
Value
Patient 09
SE_6399-W_TU2
SE_6399-W_TU1
PE_6399-W_TU2
PE_6399-W_TU1
PE_6399-W_NO
SE_6399-W_NO
SE_6399-W_NO
PE_6399-W_NO
PE_6399-W_TU1
PE_6399-W_TU2
SE_6399-W_TU1
SE_6399-W_TU2

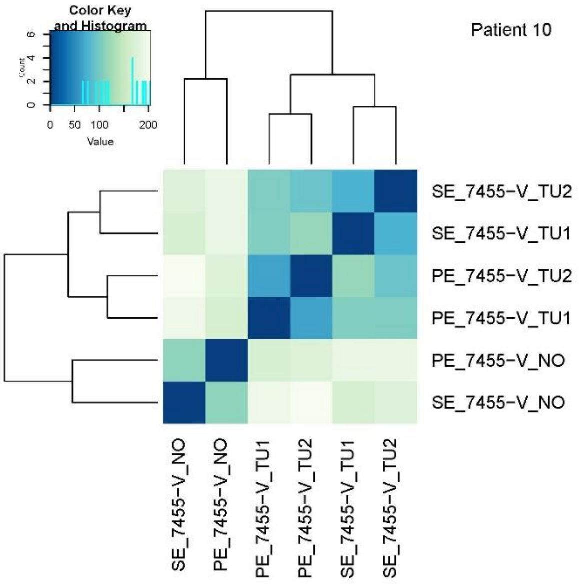
Color Key
and Histogram
Value
Patient 10
SE_7455-V_TU2
SE_7455-V_TU1
PE_7455-V_TU2
PE_7455-V_TU1
PE_7455-V_NO
SE_7455-V_NO
SE_7455-V_NO
PE_7455-V_NO
PE_7455-V_TU1
PE_7455-V_TU2
SE_7455-V_TU1
SE_7455-V_TU2

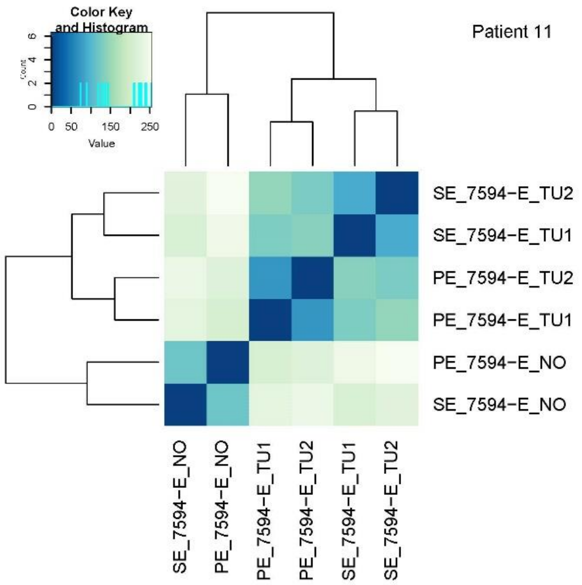
Color Key and Histogram
Count
Value
Patient 11
SE_7594−E_TU2
SE_7594−E_TU1
PE_7594−E_TU2
PE_7594−E_TU1
PE_7594−E_NO
SE_7594−E_NO
SE_7594−E_NO
PE_7594−E_NO
PE_7594−E_TU1
PE_7594−E_TU2
SE_7594−E_TU1
SE_7594−E_TU2

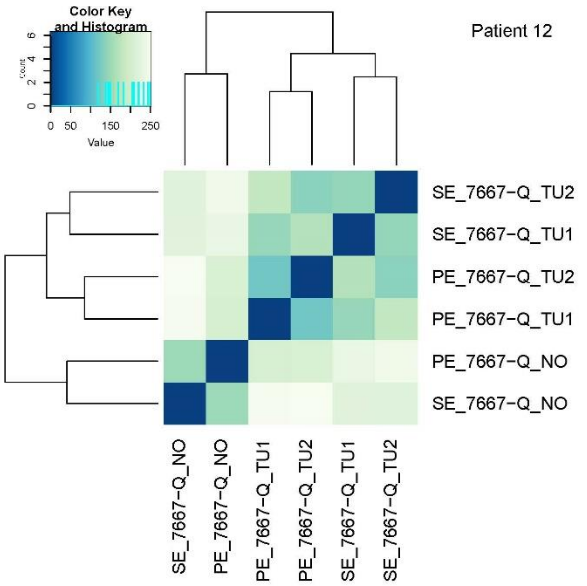
Color Key and Histogram
Count
Value
Patient 12
SE_7667−Q_TU2
SE_7667−Q_TU1
PE_7667−Q_TU2
PE_7667−Q_TU1
PE_7667−Q_NO
SE_7667−Q_NO
SE_7667−Q_NO
PE_7667−Q_NO
PE_7667−Q_TU1
PE_7667−Q_TU2
SE_7667−Q_TU1
SE_7667−Q_TU2

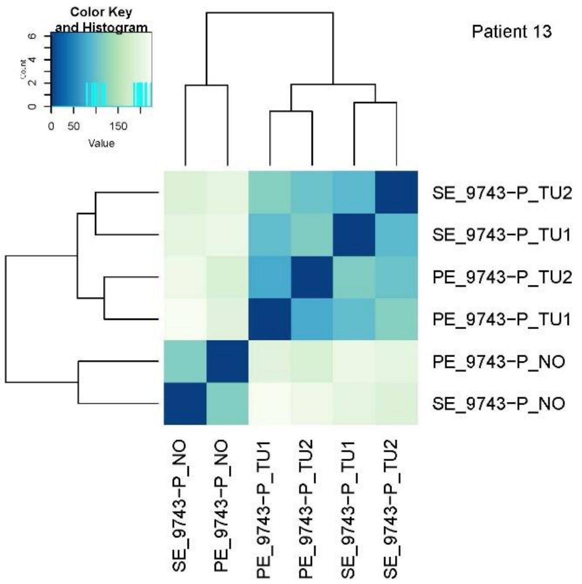
Color Key and Histogram
Count
Value
Patient 13
SE_9743−P_TU2
SE_9743−P_TU1
PE_9743−P_TU2
PE_9743−P_TU1
PE_9743−P_NO
SE_9743−P_NO
SE_9743−P_NO
PE_9743−P_NO
PE_9743−P_TU1
PE_9743−P_TU2
SE_9743−P_TU1
SE_9743−P_TU2

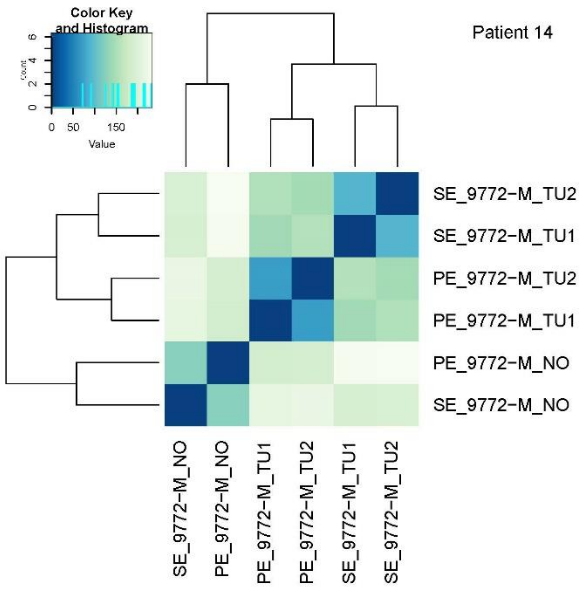
Color Key and Histogram
Count
Value
Patient 14
SE_9772−M_TU2
SE_9772−M_TU1
PE_9772−M_TU2
PE_9772−M_TU1
PE_9772−M_NO
SE_9772−M_NO
SE_9772−M_NO
PE_9772−M_NO
PE_9772−M_TU1
PE_9772−M_TU2
SE_9772−M_TU1
SE_9772−M_TU2

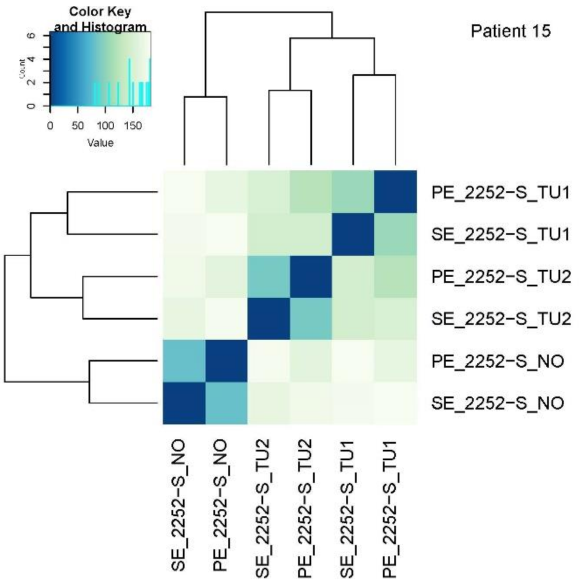
Color Key and Histogram
Count
Value
Patient 15
PE_2252−S_TU1
SE_2252−S_TU1
PE_2252−S_TU2
SE_2252−S_TU2
PE_2252−S_NO
SE_2252−S_NO
SE_2252−S_NO
PE_2252−S_NO
SE_2252−S_TU2
PE_2252−S_TU2
SE_2252−S_TU1
PE_2252−S_TU1

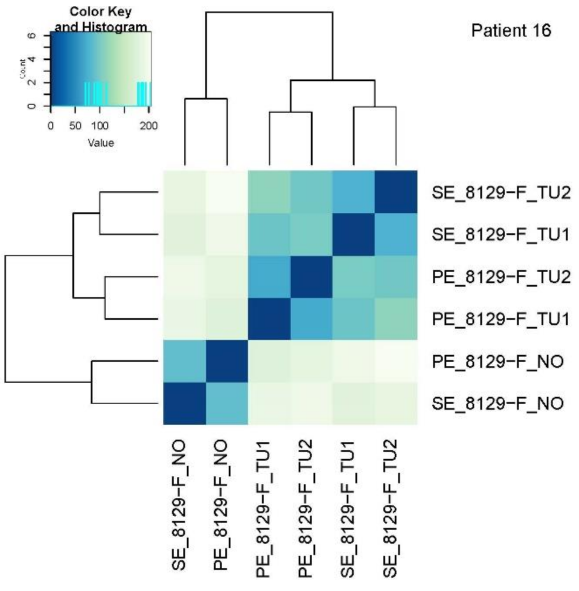
Color Key and Histogram
Count
Value
Patient 16
SE_8129−F_TU2
SE_8129−F_TU1
PE_8129−F_TU2
PE_8129−F_TU1
PE_8129−F_NO
SE_8129−F_NO
SE_8129−F_NO
PE_8129−F_NO
PE_8129−F_TU1
PE_8129−F_TU2
SE_8129−F_TU1
SE_8129−F_TU2

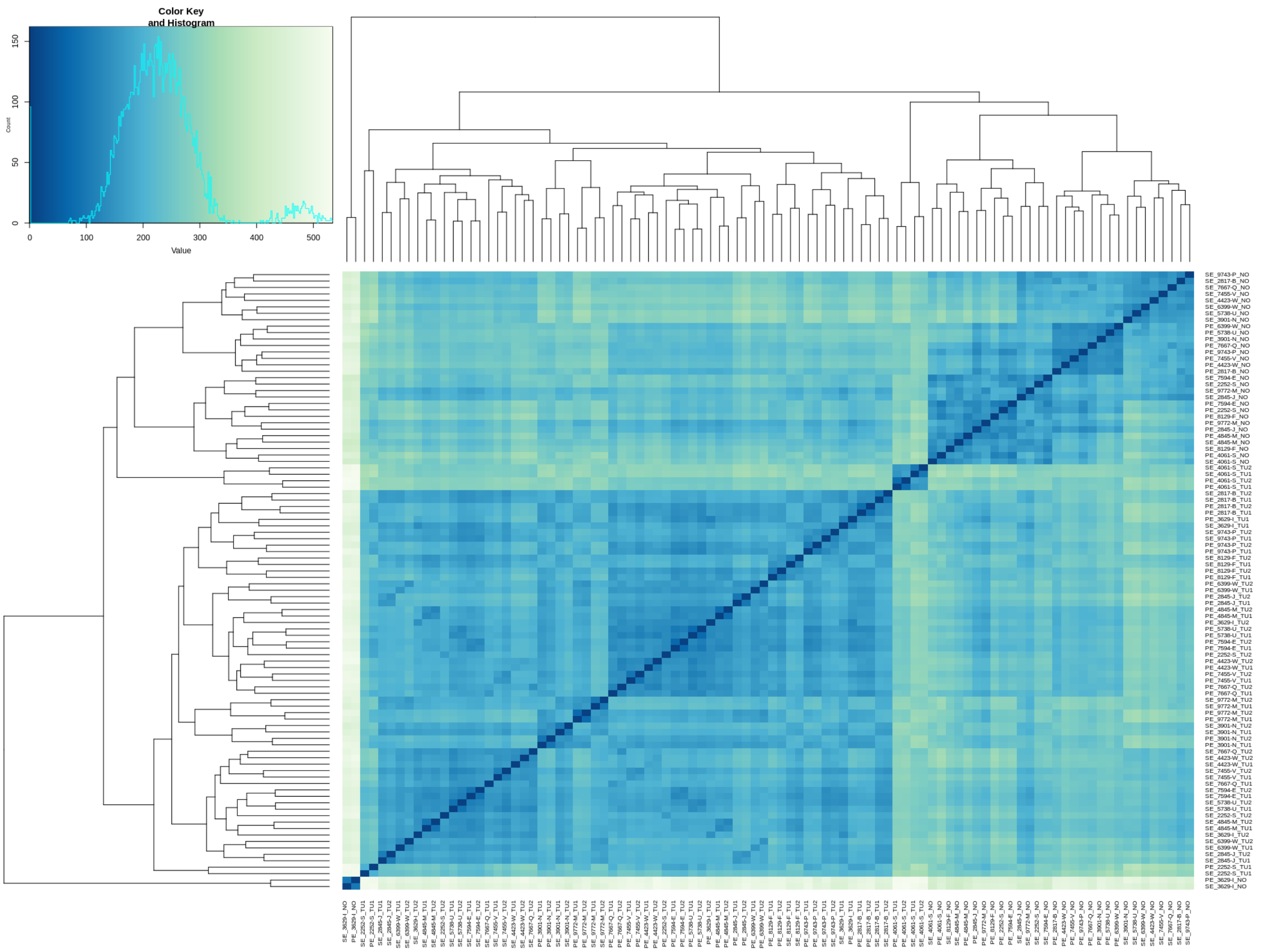
Color Key
and Histogram
Value
0
100
200
300
400
500
0
50
100
150

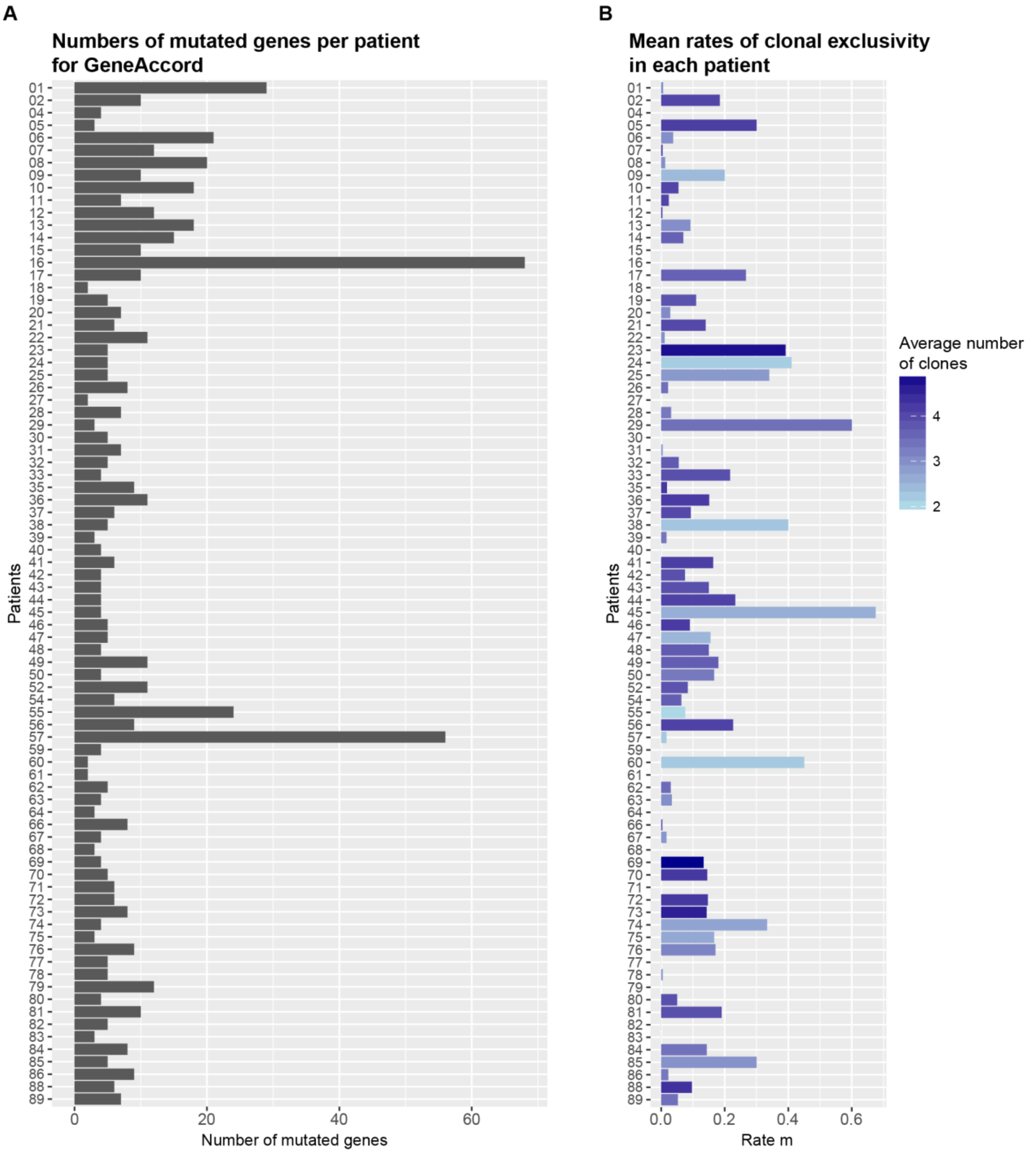


**Supplementary Figure S2. (A)** Numbers of mutated genes per patient that were used as input for GeneAccord. The mutations were assigned to clones through Cloe, and were mapped to genes afterwards. For GeneAccord, only mutations with a potential impact were considered, that is, for instance, synonymous or intronic mutations were filtered out (Methods section). Seven patients had only zero or one gene-to-clone assignment (patients 03, 34, 51, 53, 58, 65, 87), and therefore the GeneAccord analysis was done with the remaining 82 patients. **(B)** Average rates of clonal exclusivity per patient. The average rate and number of clones with at least one nonsynonymous mutation are computed from all 20 trees. The average number of clones ranges from 2.0 to 4.95. The rate of clonal exclusivity of a patient is the fraction of gene pairs that are in different branches of the phylogenetic tree. For instance, if the rate is zero, it means that the tree was linear without any branches, and that all genes assigned to the clones are in the same lineage, as is the case for patient 30.

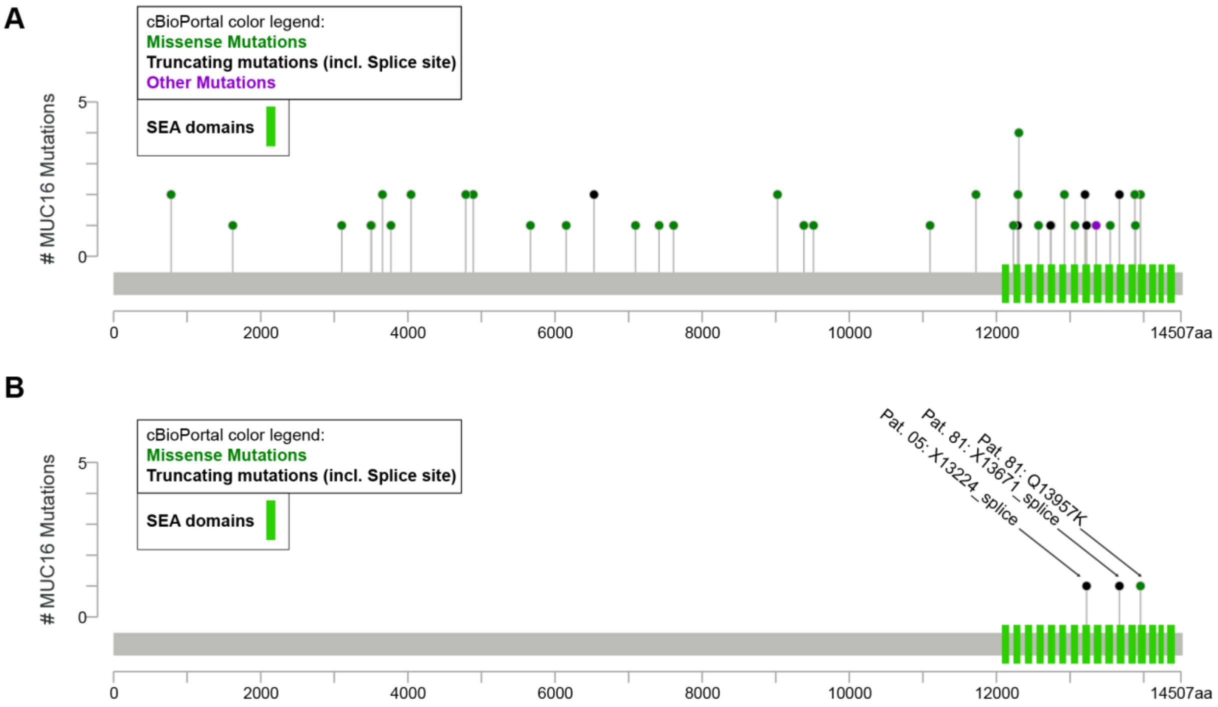


**Supplementary Figure S3. (A)** All non-synonymous and splice site mutations in *MUC16* across the samples and their location within the MUC16 amino acid sequence. The purple mutation was annotated by SnpEff as “5_prime_UTR_premature_start_codon_gain_variant”, but the cBioPortal annotated it as “Silent”. The green bars at the end of the sequence represent SEA domains of the protein. We observe that the mutations seem to preferentially affect these SEA domains. **(B)** The non-silent *MUC16* mutations in the patients 05 and 81. These were the patients with the clonally exclusive pattern of *MUC16* and *TP53*. The mutations in these patients also occur in SEA domains of the protein.

**Supplementary Table S1. The 16 gene pairs with a positive parameter delta of the clonal exclusivity test applied to the 89 ccRCC**

| Gene A | Gene B | P-value | Adjusted p-value | Mutated in (rate) | Clonally exclusive in |
|---|---|---|---|---|---|
| *MUC16* | *TP53* | < 0.01 | 0.01 | Patient 05 (0.3); Patient 81 (0.19) | Patient 05; Patient 81 |
| *BAP1* | *TP53* | 0.01 | 0.04 | Patient 30 (0); Patient 84 (0.14) | Patient 84 |
| *CACNA1S* | *MAP3K3* | 0.01 | 0.04 | Patient 16 (0); Patient 36 (0.15) | Patient 36 |
| *CSMD1* | *MAP3K3* | 0.01 | 0.04 | Patient 16 (0); Patient 36 (0.15) | Patient 36 |
| *USP36* | *CNOT1* | 0.02 | < 0.05 | Patient 16 (0); Patient 49 (0.18) | Patient 49 |
| *PBRM1* | *BAP1* | 0.02 | 0.05 | Patient 30 (0); Patient 35 (0.02); Patient 84 (0.14) | Patient 84 |
| *MUC16* | *TRO* | 0.03 | 0.05 | Patient 26 (0.02); Patient 56 (0.23) | Patient 56 |
| *DNAH9* | *MUC16* | 0.04 | 0.05 | Patient 06 (0.04); Patient 56 (0.23) | Patient 56 |
| *DNAH9* | *WNT5B* | 0.04 | 0.05 | Patient 06 (0.04); Patient 17 (0.27) | Patient 17 |
| *DNAH9* | *UNC79* | 0.04 | 0.05 | Patient 06 (0.04); Patient 17 (0.27) | Patient 17 |
| *WNT5B* | *RARA* | 0.04 | 0.05 | Patient 06 (0.04); Patient 17 (0.27) | Patient 17 |
| *RARA* | *UNC79* | 0.04 | 0.05 | Patient 06 (0.04); Patient 17 (0.27) | Patient 17 |
| *PBRM1* | *CSMD3* | 0.04 | 0.05 | Patient 73 (0.14); Patient 84 (0.14) | Patient 84 |
| *CSMD3* | *ARHGAP35* | 0.05 | 0.06 | Patient 02 (0.18); Patient 84 (0.14) | Patient 02 |
| *USP36* | *YLPM1* | 0.06 | 0.06 | Patient 16 (0); Patient 45 (0.68) | Patient 45 |
| *SETD2* | *PIK3CA* | 0.07 | 0.07 | Patient 10 (0.05); Patient 74 (0.33); Patient 82 (0) | Patient 74 |

The 16 gene pairs from the panel sequencing data set whose estimate of the parameter delta is greater zero. The column ‘Mutated in (rate)’ lists the patients in which the pair was mutated in. The average rate of clonal exclusivity of the respective patient is shown in parenthesis. The column ‘Clonally exclusive in’ lists the patients in which the pair was also clonally exclusive in the majority of the trees.

**Supplementary Table S2. The variant categories from Fig. 3 and the corresponding annotations from SnpEff**

| Variant category in Fig. 3 | SnpEff annotation |
|---|---|
| Start lost | start_lost&conservative_inframe_deletion, start_lost |
| Stop lost | stop_lost, frameshift_variant&stop_lost |
| Stop gained | stop_gained, stop_gained&disruptive_inframe_deletion, stop_gained&splice_region_variant |
| Start gained | 5_prime_UTR_premature_start_codon_gain_variant |
| Frameshift indel | frameshift_variant, frameshift_variant&splice_region_variant, frameshift_variant&splice_donor_variant&splice_region_variant&intron_variant |
| Inframe indel | disruptive_inframe_deletion, disruptive_inframe_insertion, inframe_insertion, inframe_deletion, conservative_inframe_insertion, conservative_inframe_deletion |
| Missense | missense_variant&splice_region_variant, missense_variant |
| Splice site | splice_donor_variant&splice_region_variant&3_prime_UTR_variant&intron_variant, splice_region_variant, splice_region_variant&synonymous_variant splice_region_variant&non_coding_exon_variant, splice_region_variant&intron_variant, splice_region_variant&non_coding_transcript_exon_variant, splice_acceptor_variant&intron_variant, splice_donor_variant&intron_variant |
| Protein interacion loci | structural_interaction_variant, protein_protein_contact |
| 5 prime UTR | 5_prime_UTR_variant, 5_prime_UTR_truncation&exon_loss_variant |
| 3 prime UTR | 3_prime_UTR_variant |
| Synonymous | synonymous_variant |
| Ignored | intron_variant |
| Sequence feature | sequence_feature |
| Ignored | non_coding_exon_variant, non_coding_transcript_variant, non_coding_transcript_exon_variant |
| Ignored | intergenic_region, upstream_gene_variant, downstream_gene_variant, TF_binding_site_variant(no gene associated to this annotation; it is a "motif annotation"), intragenic_variant |

The variants were annotated to genes with the program SnpEff (2). It concatenates annotations with '&' in case several effects are true for the same mutation and transcript. The same gene can be hit by several mutations, or the same mutation can have several annotations for different transcripts. Therefore, in order to visualize the mutations in a heatmap like in Fig. 3, an order of importance needed to be defined, which is the following starting with highest priority: 'Start lost', 'Stop lost', 'Stop gained', 'Start gained', 'Frameshift indel', 'Inframe indel', 'Missense', 'Splice site', 'Protein interaction loci', '5 prime UTR', '3 prime UTR', 'Synonymous', 'Sequence feature'. This means that, if a gene was hit by two mutations, where one is synonymous and the other one is a missense mutation, only the missense mutation will be shown in the heatmap.

**Supplementary Table S3. The ten most striking pathway pairs with a positive parameter delta of the clonal exclusivity test applied to the 16 ccRCC on the pathway level**

| Pathway A | Pathway B | P-value | Adjusted p-value | Affected in (rate) | Clonally exclusive in |
|---|---|---|---|---|---|
| Defective B3GALTL causes Peters-plus syndrome (PpS) | Major pathway of rRNA processing in the nucleolus and cytosol | 0 | 0 | Patient 08 (0.012); Patient 14 (0.008) | Patient 08; Patient 14 |
| O-glycosylation of TSR domain-containing proteins | Major pathway of rRNA processing in the nucleolus and cytosol | 0 | 0 | Patient 08 (0.012); Patient 14 (0.008) | Patient 08; Patient 14 |
| Cobalamin (Cbl, vitamin B12) transport and metabolism | PRDM12 | 0 | 0.02 | Patient 04 (0.003); Patient 15 (0.008) | Patient 15 |
| COPI-dependent Golgi-to-ER retrograde traffic | Major pathway of rRNA processing in the nucleolus and cytosol | 0.001 | 0.026 | Patient 03 (0.029); Patient 14 (0.008) | Patient 14 |
| Kinesins | Major pathway of rRNA processing in the nucleolus and cytosol | 0.001 | 0.026 | Patient 03 (0.029); Patient 14 (0.008) | Patient 14 |
| Major pathway of rRNA processing in the nucleolus and cytosol | MHC class II antigen presentation | 0.001 | 0.026 | Patient 03 (0.029); Patient 14 (0.008) | Patient 14 |
| Activation of Matrix Metalloproteinases | Immunoregulatory interactions between a Lymphoid and a non-Lymphoid cell | 0.001 | 0.026 | Patient 01 (0.015); Patient 10 (0.017); Patient 15 (0.008) | Patient 15 |
| Ion homeostasis | NCAM1 interactions | 0.002 | 0.026 | Patient 02 (0.003); Patient 06 (0.047); Patient 15 (0.008) | Patient 02 |
| E3 ubiquitin ligases ubiquitinate target proteins | MUC2 | 0.002 | 0.026 | Patient 06 (0.047); Patient 13 (0.065) | Patient 13 |
| Laminin interactions | MUC2 | 0.005 | 0.026 | Patient 06 (0.047); Patient 13 (0.065) | Patient 13 |

The ten most striking pathway pairs from the WES data set whose estimate of the parameter delta is greater zero. The column ‘Affected in (rate)’ lists the patients in which the pair was affected in. The average rate of clonal exclusivity of the respective patient is shown in parenthesis. The column ‘Clonally exclusive in’ lists the patients in which the pair was also clonally exclusive in the majority of the tree inference runs. In case a gene could not be mapped to a pathway, as is the case for PRDM12 and MUC2, the gene identifiers were retained.
